# Diapause is not selected as a bet-hedging strategy in insects: a meta-analysis of reaction norm shapes

**DOI:** 10.1101/752881

**Authors:** Jens Joschinski, Dries Bonte

## Abstract

Many organisms escape from lethal climatological conditions by entering a resistant resting stage called diapause, which needs to be optimally timed with seasonal change. As climate change exerts selection pressure on phenology, the evolution of mean diapause timing, but also of phenotypic plasticity and bet-hedging strategies is expected. Especially the latter as a strategy to cope with unpredictability is little considered in the context of climate change.

Contemporary patterns of phenological strategies across a geographic range may provide information about their evolvability. We thus extracted 458 diapause reaction norms from 60 studies. First, we correlated mean diapause timing with mean winter onset. Then we partitioned the reaction norm variance into a temporal component (phenotypic plasticity) and among-offspring variance (diversified bet-hedging) and correlated this variance composition with predictability of winter onset. Mean diapause timing correlated reasonably well with mean winter onset, except for populations at high latitudes, which apparently failed to track early onsets. Variance among offspring was, however, limited and correlated only weakly with environmental predictability, indicating little scope for bet-hedging. The apparent lack of phenological bet-hedging strategies may pose a risk in a less predictable climate, but we also highlight the need for more data on alternative strategies.

## Introduction

Anthropogenic greenhouse gas emissions change the environment, most notably temperatures, at an unprecedented rate [1], and the majority of species face extinction risks from climate change [2]. One of the most commonly observed responses to climate change is a shift in phenology, i.e. in the seasonal timing of an organism [3]. Changes in tree leaf-out [4] and bird egg-laying dates [5] in spring are among the most famous examples of phenology shifts, but shifts in timing have been documented across nearly the whole tree of life (e.g. cyanobacteria [6], fungi [7], cnidarians [8], insects [9]). Phenological shifts that match an organism’s life cycle with novel conditions can generally be expected to provide fitness benefits [10] (but see [11]), but there is increasing doubt that such phenological shifts will remain sufficient in a rapidly changing climate [12]. Hence it is essential to infer the evolutionary potential of phenological strategies.

Phenology is subject to multiple selection pressures, rendering any predictions on its evolvability not straightforward. For example, variation in the extent of phenology shifts among interacting species may create mismatches [10,13], thus potentially selecting against phenology shifts. Moreover, novel correlations of temperature and day length may impose physiological constraints, such as day length limitations for diurnal animals [14–16] and plants [17] (but see [18]) – relying on mistimed developmental cues may then constitute an evolutionary trap [19,20]. Therefore it is not clear whether a complex trait such as phenology can evolve an optimal response to changing local conditions. A longitudinal analysis about adaptation of phenology in the past, across species and habitats, may however identify potential evolutionary constraints that could also impede adaptation to a changing climate.

Rises in mean temperatures are not the only potential cause of climate-change induced biodiversity loss - increasing climate variability imposes further extinction risk [21]. Therefore, the concerted evolution of mean phenology and risk-reduction strategies will be required. There are three general strategies by which organisms can cope with changing environments [22–24]: evolution of the mean, phenotypic plasticity, and bet-hedging (avoidance of fitness variance; see also [25] for examples). The latter consists of strategies to avoid risk (conservative bet-hedging) and of strategies to spread the risk among one’s offspring (diversified bet-hedging). These strategies are intricately related [20,23,26,27], which make examining their simultaneous evolution a daunting task. However, for polyphenisms with only two outcomes, such as the decision to overwinter or to germinate, the strategies can be conveniently separated by correlating the reaction norm shape with climatic conditions [25]: the variance composition, i.e. the ratio of variance among environments vs. variance among the offspring, determines the degree of diversified bet-hedging and plasticity, while the reaction norm mean determines the distinction among conservative bet-hedging and arithmetic mean optimization (see methods, Fig. 1).

**Fig. 1.**
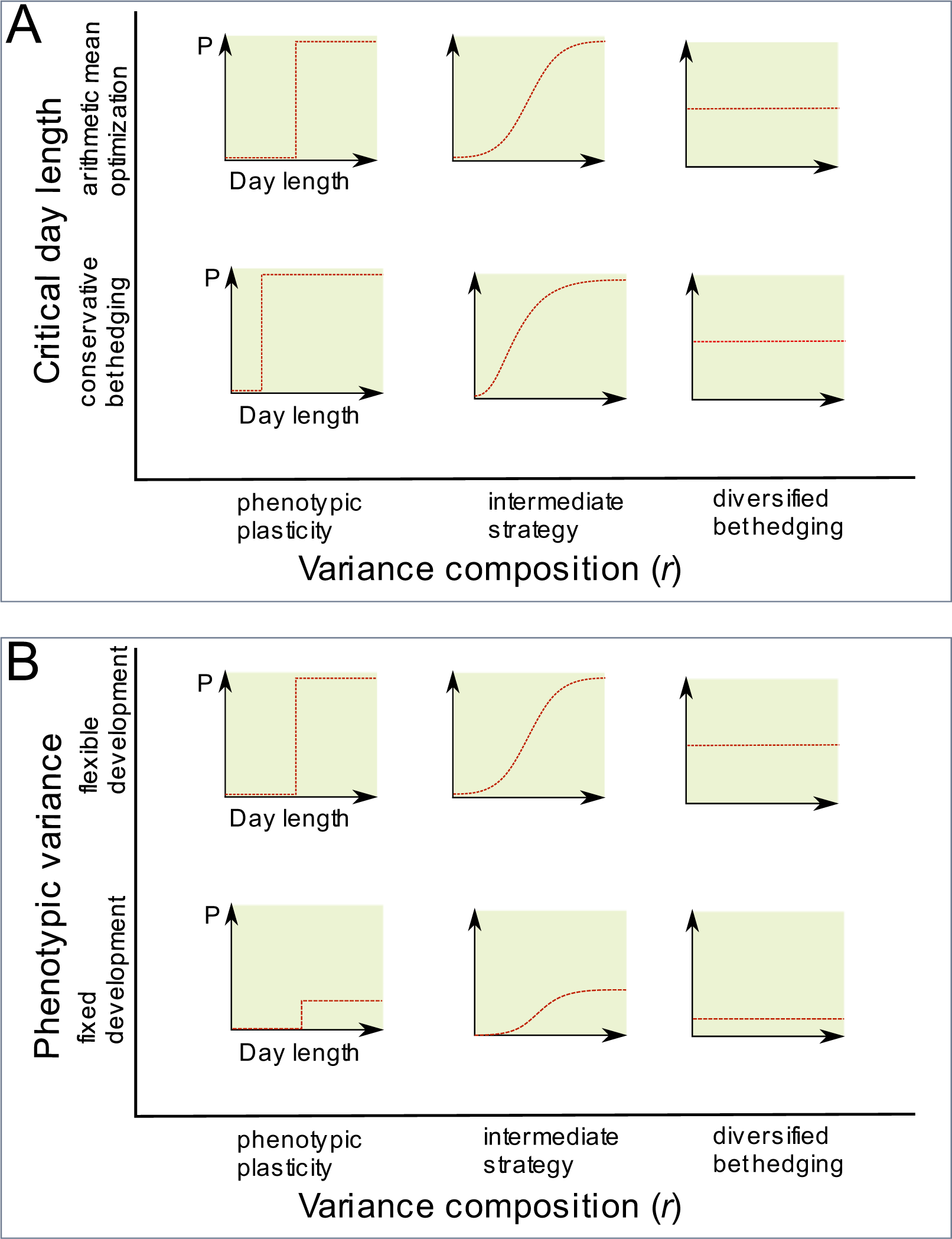
Relationship of evolutionary strategies with reaction norm properties. Panel A shows a series of dichotomous reaction norms, in which the proportion of phenotypes (P) can range from zero to 1. The decision to switch phenotypes can be expressed by a steep logistic curve (upper left), but reaction norms can also divert in various ways from this step function: By changes in the ratio of the variance components among 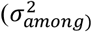 and within 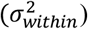 environments (x-axis), and by a shift in the mean frequency, or inflection point (y-axis). Moreover, the sum of the variance components may also vary, as reaction norms may be canalized (Panel B). For details on this framework, see Joschinski & Bonte (2019). 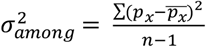;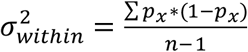.

While many studies that research adaptation to climate change focus on spring phenology, changes in autumn phenology remain relatively understudied [28]. The induction of winter resting stages is generally governed by day length [29,30], as seasonal variation in day length and in energy transfer from the sun share the same cause (axial tilt of the Earth) and are hence correlated. Although there are exceptions in which temperature [31] or other cues [32] play a major role, photoperiodism remains the best studied phenological trait to date [33]. Insect winter diapause, a resting stage to overwinter, is a polyphenism that is particularly well-studied [34–36], and there is ample high-quality data under laboratory conditions available. More than 50 years ago it has been reported that the critical day length, i.e. the inflection point of diapause reaction norms increases with latitude, at a rate of 60 – 90 minutes in day length change per 5°N [34].We collected 458 facultative diapause reaction norms from laboratory experiments (60 studies; Supplementary material S1), derived their critical day length (which determines mean diapause timing) and the variance composition, and then correlated them with mean winter onset and winter predictability as derived from climate data. First, we estimated by how much the critical day length changes with latitude, thereby validating earlier case studies based on less robust data [34]. Then we used these data to seek whether theoretical predictions on bet hedging and plasticity hold for insect diapause as one of the most critical traits for insect fitness. More specifically we tested whether

1. Mean diapause correlates with mean winter onset (*arithmetic mean optimization*);
2. The variance composition of reaction norms correlates with environmental predictability (*phenotypic plasticity / diversified bet-hedging*); and
3. Deviation from optimal mean timing towards early diapause correlates with environmental predictability (*conservative bet-hedging*)

## Methods

### Rationale and effect size calculation

A full description about the relationship of polyphenic reaction norm shapes with evolutionary strategies can be found elsewhere [25]. In short, we see phenotypic plasticity and diversified bet-hedging as opposite ends on a continuum of reaction norm shapes. One extreme is a steep (“plastic”) reaction norm that creates maximal variance among environments (Fig. 1A, upper left). We define this variance among environments as the squared standard deviation of probabilities,

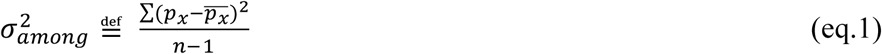

with *p*_*x*_ being the probability of diapause in environment *x*, and *n* being the number of environments.

The other extreme ensures equal production of both phenotypes under all environmental conditions, and is thus flat at the 50% level (Fig. 1A, upper right). This shape maximizes the variance among the offspring, i.e. the variance within rather than among environments. This variance component can be described as a series of Bernoulli draws along an environmental gradient (day length treatments):

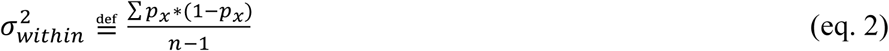

Between those extremes lies a continuum of reaction norm shapes (Fig. 1A, middle column) that can be described by the ratio of the variances. This *variance composition* is thus:

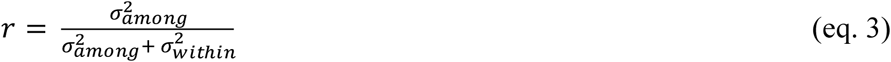

In summary, *r* determines the variance in offspring phenotypes, and leads to a continuum from diversified bet-hedging to phenotypic plasticity. The mean of the offspring distribution may, however, also vary, governing a continuum from arithmetic mean optimization to conservative bet-hedging. For logistic reaction norms, the mean of the offspring distribution is determined by the inflection point (Fig. 1A, lower row), which is also called critical day length in diapause reaction norms [34].

For the sake of completeness, we also described a third dimension of reaction norm shapes [25], which regards the phenotypic variance (Fig. 1B). Phenotypic variance can be described as the sum of the two variance components. A reaction norm that is flat at 0 % or 100% diapause (i.e. canalized, or environment-independent) exhibits neither variance among (plasticity) nor within environments (diversified bet-hedging), and represents obligate development or obligate diapause. This reaction norm dimension was, however, not further studied in our meta-analysis, because our literature search was implicitly biased against canalized reaction norms (see search criteria and discussion). Hence the two effect sizes considered in our meta-analysis were the variance composition (eq. 3) and the inflection point. In classical diapause experiments, insects have been subjected to multiple day lengths and the percentage of diapause induction was recorded [34–36], the reaction norm is therefore only approximated with discrete day lengths. To arrive at a continuous reaction norm, we modelled reaction norm shape via four parameters:

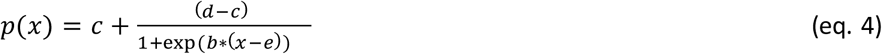

In this equation *p(x)* is the frequency of diapausing individuals under day *x*. *e* is the inflection point of the curve, i.e. the critical day length, and hence directly represents the axis *mean*. *c* and *d* indicate the lower and upper diapause threshold, and *b* is the slope of the curve.

### Empirical data

#### Literature search

We concentrated on studies that measure photoperiodic responses of terrestrial arthropods, though invertebrates with a larval stage in shallow water (e.g. mosquitoes) were also included. We only used studies with at least four photoperiods treatments as this is the minimal requirement to construct 4-parameter logistic growth curves (eq. 4). We did not restrict our analysis to any geographic location or publication language, but we selected only studies with at least three populations. We conducted two independent literature searches in the Web of Science core collection, the Russian Science Citation Index, the KCI Korean Journal Database and the SciELO database (Fig. 2) on 10.12.2018. First we limited the search terms to:

**Fig. 2.**
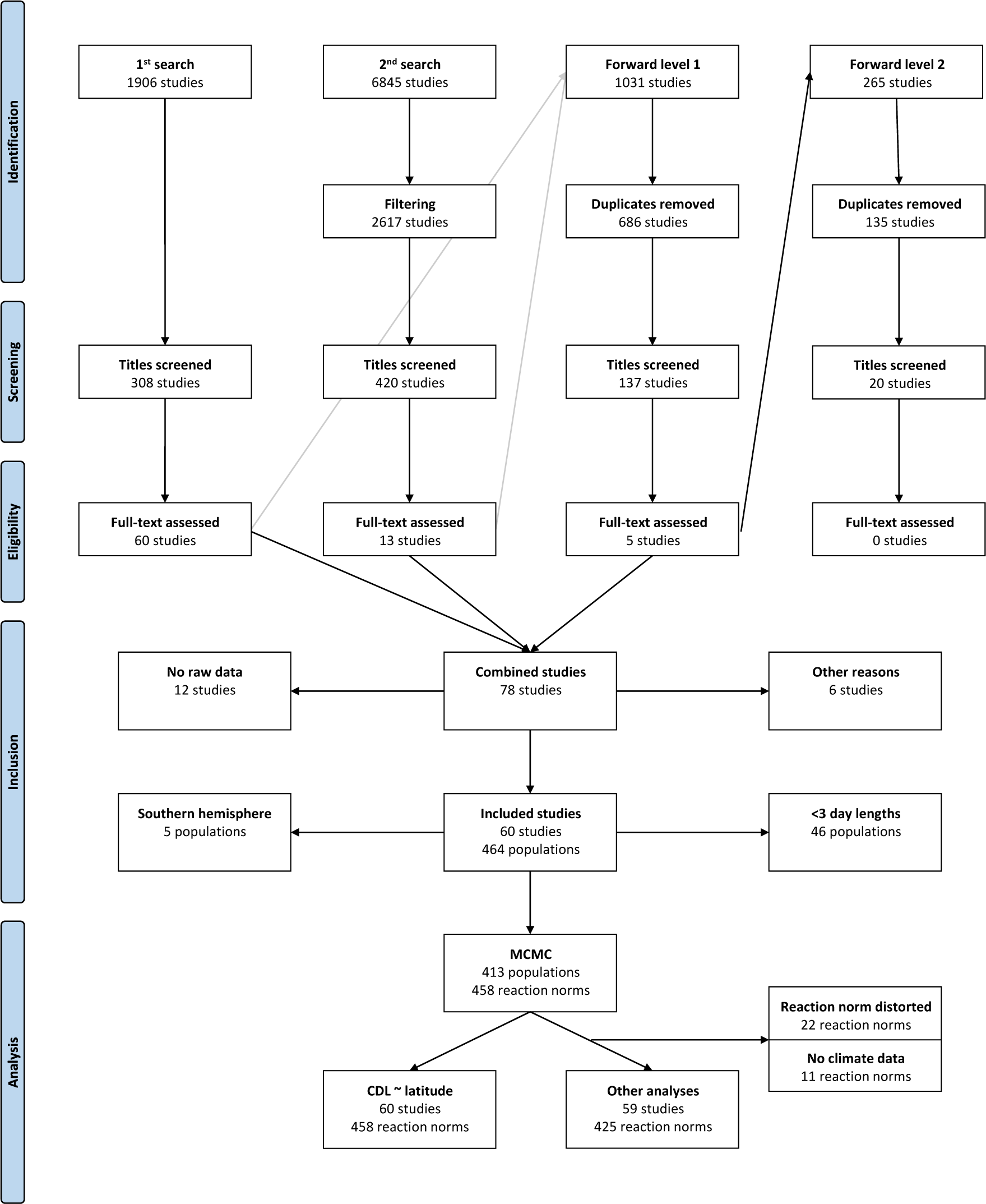
Prisma diagram describing the literature search and sample sizes for statistical analysis of diapause reaction norm shapes.

*TS = ((photoperiodic AND (geogr* OR range)) OR (photoperiod* AND latitud*) OR (photoperiod* AND longitud*))*

We found 1906 references in the first search, of which we judged 308 potentially relevant, and 60 met all eligibility criteria. Secondly, we used a wider range of search terms:

*TS = (("day length" OR photoperiod* OR diapaus* OR hibern* OR dorman*) AND (geogr* OR "range" OR latitud* OR longitud* OR cline$ OR clinal))*

We excluded all articles that were found in the first search, as well as all review articles, retractions and corrections. We then filtered the 6845 results by research area and invertebrate related terms (Supplementary material S2). 2617 articles remained, with 420 potentially relevant and 13 eligible articles. We did a forward-citation search over all databases on the 73 eligible articles of both searches on 12.12.2019 and found 686 new references, which included 137 potential and 5 eligible articles. A second forward-citation search on these five articles on 12.12.2019 brought 135 new articles, but none were relevant. Altogether there were 78 eligible references.

#### Inclusion criteria

12 articles were excluded because they were not accompanied by raw data, tables or figures that allowed further analysis, and the authors were deceased, did no longer have the raw data or did not respond to our emails. We further removed six articles that were otherwise not usable, so 60 studies with 464 populations remained. We removed 46 individual populations with less than four day length measurements from these studies, as well as five populations from the southern hemisphere, so 413 populations remained. Because some studies reported reaction norms for multiple lines from the same population, there were 458 reaction norms available, and these 458 reaction norms consisted of 3092 individual data points.

#### Data extraction

The reaction norms in 51 of the 60 studies were presented as figures. In these cases we saved the figure and extracted the data with WebPlotDigitizer Version 3.12 [37]. Where necessary, the day length was then rounded or corrected to match the description in materials and methods of the respective study. Y-values that were slightly above 100% or below 0% were set to 100% and 0% respectively.

Detailed information on numbers of individuals per day length estimate were rarely available (100 reaction norms), as numbers were either given as population-level means (30 reaction norms), as global average or range (300 reaction norms), or missed entirely (33 reaction norms). We wish to emphasize that a lack of detailed information should not be confused with an unweighted (“vote-count”) meta-analysis, because the sample size (day lengths per population) was always known. Rather, the missing information occurred on a lower level (points within population) than the level of replication (population). Where the data was provided, we recorded it for later weighing of the data points.

#### Calculation of mean and variance composition

The published reaction norms reported the change of diapause percentages with day length. Day length depends, however, on latitude, and thus is not a direct indicator of phenology, so we converted day lengths into ordinal days by using the reported latitude of the sampling location and the *daylength* function from the package *geosphere* [38]. 743 of the 3092 day length treatments were outside naturally occurring day lengths. We assumed that these artificial day lengths represent diapause incidence at midsummer and midwinter, respectively, but removed 22 reaction norms that became severely distorted by this assumption. All further analysis except the correlation of critical photoperiod with latitude are based on the converted reaction norms.

To derive the effect sizes (means and variance composition) we modelled continuous reaction norms with a Markov chain Monte Carlo method [39] according to eq. 4 (Supplementary material S3).

### Climate data

We used land surface temperature data from the Global Historical Climatology Network GHCN-Daily [40,41]. We extracted daily minimum and maximum temperatures from ~34,000 climate stations and then calculated daily mean temperature as the average of the extremes. After cleaning the data to stations in the northern hemisphere and with at least 3 years of data with 180 temperature records, the data consisted of 10,991,727 months (3-244 years) in 26,804 climate stations.

To estimate winter onset in each year and station, we identified cold days with average temperatures below 10°C. We then determined winter onset as the fifth cold day after midsummer. Years in which winter did not arrive according to this definition were excluded, and stations with less than 3 years with winter onset removed. We calculated a weighted mean winter onset and a frequency weighed standard deviation of winter onset to account for differences in reliability (days with eligible data) across years. Stations with standard deviations in winter onset above 30 (4.2% of all stations) were then also deemed unreliable and removed. We obtained 24,266 estimates of mean winter onset, day length at winter onset and winter predictability in the northern hemisphere.

Initial data handling was performed with a perl script, whereas all further analysis was conducted in R version 3.4.3 [42], using R base functions and convenience functions [43–51].

#### Merging with empirical data

To combine climate data and study site locations, we averaged the climate estimates from the 5 closest stations within a 5° radius (weighted by 1/Euclidian distance). When the coordinates were not directly provided in the study, we used the coordinates of the quoted town or area. Town and area coordinates were made available by the WikiProject Geographical coordinates (https://en.wikipedia.org/wiki/Wikipedia:WikiProject_Geographical_coordinates) and the Geohack tool (https://www.mediawiki.org/wiki/GeoHack). 11 populations did not have any climate station nearby and were only used for correlations with latitude, but not in further analyses.

### Analysis

We used linear mixed-effects models with a nested random structure [52] to correlate the reaction norm properties with climate variables. The random effects were nested on five levels (order/genus/species/study/population), but we simplified the random structure to order/species/population, ignoring both study ID and genus. Study ID was disregarded because most species were only represented by a single study, and the 12 species that were represented by multiple studies usually contained the same first or lead authors and applied the same methods (Supplementary material S1). Genus was disregarded because there were either only very few genera per order available (e.g. Diptera), or all species within an order were placed in different genera (Lepidoptera, Supplementary material S1). We weighed the reaction norm estimates by the inverse of the variance (credible interval ranges, divided by 2*1.96 and squared), but truncated the intervals at the lower limit to a biologically meaningful value to prevent some estimates from obtaining nearly infinite weight.

We performed the following statistical models: first, we correlated critical photoperiod (the inflection point of the day length reaction norm) with latitude. This correlation systematically tests Danilevskii’s observation that the critical photoperiod changes by 60-90 minutes per 5° latitude [34]. Next, we converted the critical photoperiod into ordinal days. The mapping of critical photoperiod to day of the year is latitude – dependent, so mean diapause timing cannot be expressed by critical photoperiod alone (see discussion). We correlated this converted diapause timing with latitude. Thirdly, we correlated mean diapause timing (converted as before) with mean winter onset as derived from climate data. This correlation tested whether diapause is timed such that the arithmetic mean is optimized (but see limitations section). To test for potential adaptive phenotypic plasticity and diversified bet-hedging, we then correlated the variance composition determining the level of diversified bet-hedging to phenotypic plasticity (eq. 3) with winter predictability. Lastly, we tested for conservative bet-hedging, i.e. whether populations from unpredictable climates diapause earlier than expected based on mean winter onset. To do so, we correlated the residuals of our third model (mean diapause ~mean winter onset) with winter predictability.

We truncated the credible interval minimum (see above) to 10 minutes in the day length reaction norms, to 1 week in the ordinal day reaction norms, and to 5% in the remaining models.

The *phenotypic variance* (obligate vs. facultative diapause) was not further studied, because we explicitly searched for day length reaction norms, thereby excluding species that do not rely on day length for their phenology.

We used the full dataset (458 reaction norms) for the day length reaction norm model but removed all reaction norms that were not convertible into ordinal days or had no nearby climate stations (425 remaining) for all other models.

We assumed a gaussian distribution for all models, though we logit-transformed variance ratios prior to analysis. For all models we report partial R² values, calculated as proportion of variance reduction at each random level, 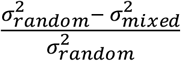. This is an extension of a commonly used pseudo-R^2^ value [53]. In addition, we provide likelihood ratio test statistics (model with and without fixed effect). Model 3 was conducted without the nested random terms, because their effect was already accounted for by model 1.

#### Sensitivity of climate predictions to temperature threshold

Arthropod thermal requirements vary among species, and our use of a 10°C temperature threshold was an arbitrary decision. It resulted in a global median winter onset around Oct 11, which is within the range of commonly reported phenological windows and threshold values [54,55]. To explore the sensitivity of our meta-analysis to the arbitrary threshold, we systematically varied it between 0 and 15°C, and calculated the R² profiles of models 1 and 2.

## Results

The climate data indicated that the timing of winter onset (by which we mean the onset of cold days in autumn) was consistently earlier at higher latitudes and altitudes (Fig. 3A). While this relationship between mean winter onset and latitude was nearly linear (not shown), the day length which corresponds to mean winter onset increased exponentially with latitude, as high latitudes featured both earlier winter onset and longer autumn days. Between 21 and 69°N we predicted day length at winter onset to decline by 46.58 minutes per 5° latitude (R² = 0.54; Supplementary material S4).

**Fig. 3.**
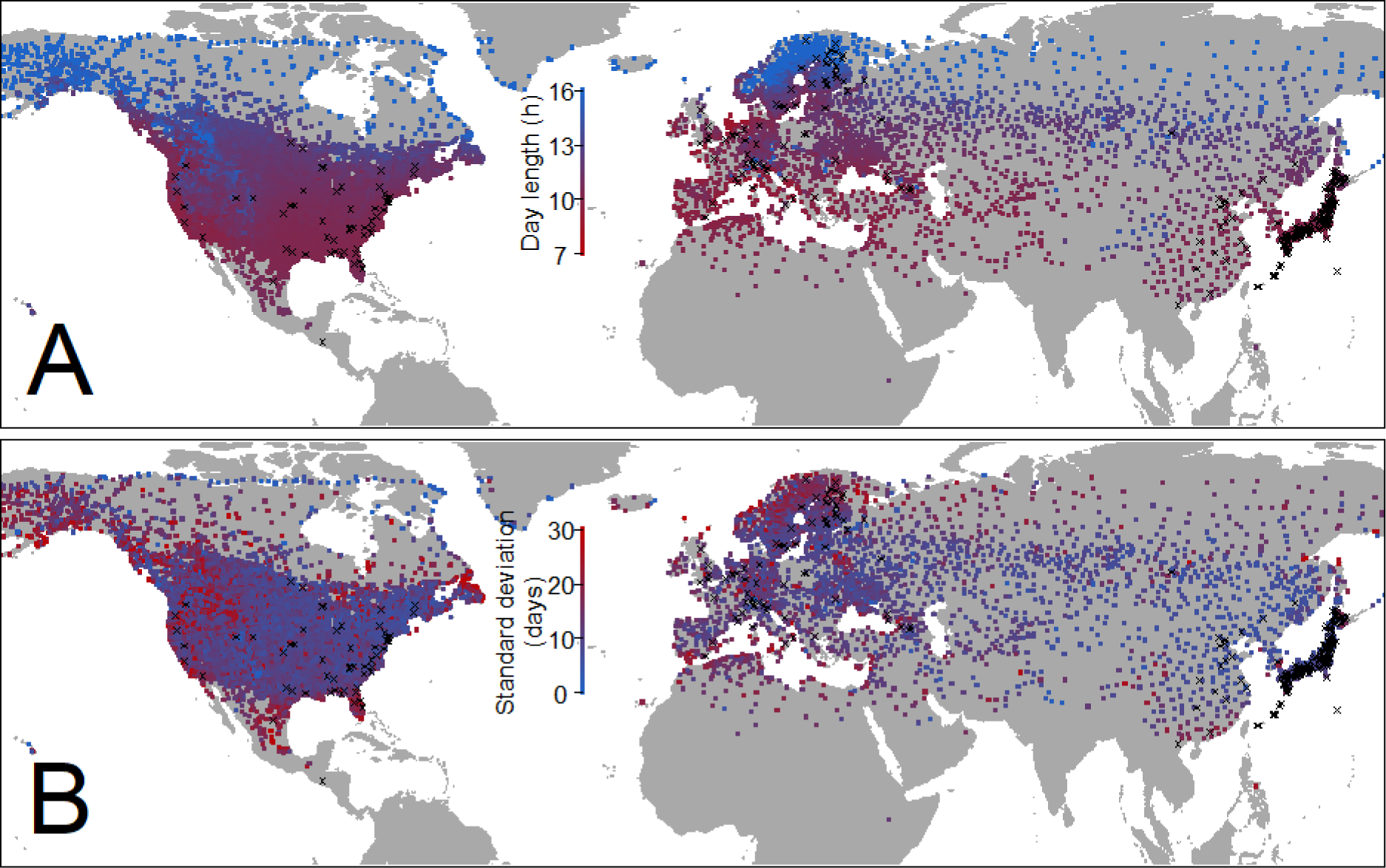
Winter onset calculation based on GHCN-daily climate data. **A**: Mean day length at winter onset, **B**: Standard deviation in winter onset. Black crosses: sampling locations of empirical studies.

For our meta-analysis we found 60 studies that met all inclusion criteria. These 60 studies featured 458 reaction norms from 8 invertebrate orders (insects and mites), though we note strong geographical and phylogenetic clustering of the data. For example, more than half of the data stem from Japan, most studies on Dipterans were conducted on Drosophila species, and most high-latitude populations are Dipterans (Supplementary Material S1). The critical day lengths (median day length that induces diapause) of the reaction norms correlated with latitude, showing a linear increase by 48.54 ± 1.89 min per 5 ° N (Fig. 4A; 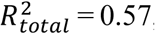, 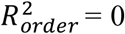, 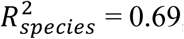, 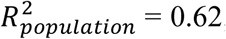; LRT ratio = 399.1, p < 0.0001). Upon conversion to ordinal days (day of the year), the inflection points correlated less strongly with latitude (Supplementary material S5; 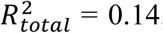, 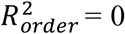, 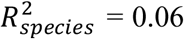, 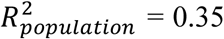 LRT ratio = 141.8, p < 0.0001) and mean winter onset (Fig. 4B; 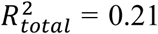, 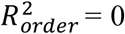, 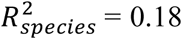, 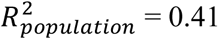, LRT ratio = 175.7, p < 0.0001). The correlation was most notably disturbed by Drosophilid populations from climates with an early winter onset (high latitudes), which diapaused later than expected based on climate data, and additionally showed little spread in mean diapause timing.

**Fig. 4.**
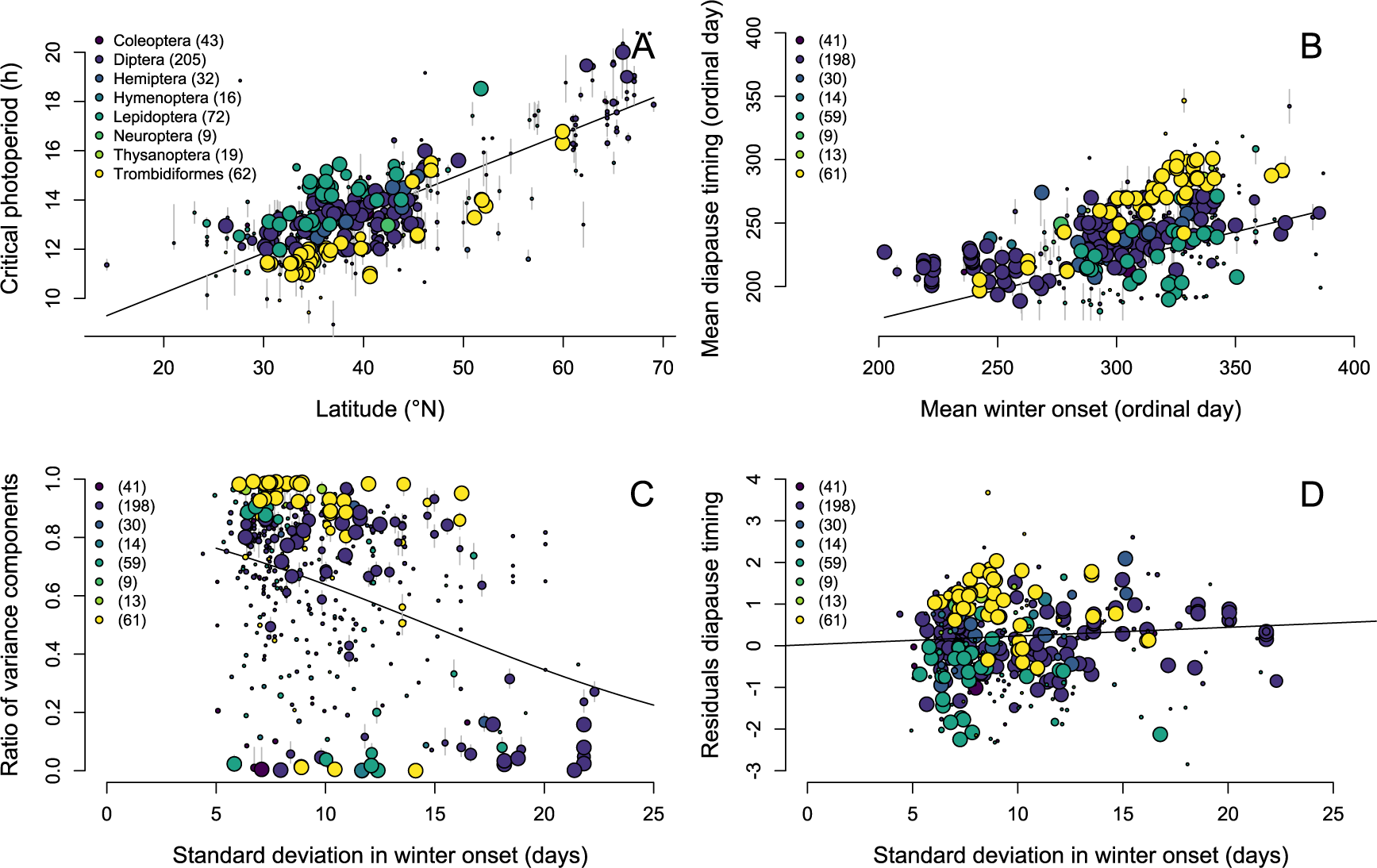
Correlation of reaction norm properties with climate variables. **A:** Critical photoperiod from primary studies versus latitude; **B:** Correlation of mean diapause timing with mean winter onset; **C:** Variance composition versus day length predictability. Ratios of 0 indicate the potential for pure bet-hedging strategies, ratios of 1 equal purely plastic strategies; **D:** Residual deviation from diapause timing against winter predictability (conservative bet-hedging). Each data point represents 1 reaction norm (458 in total), size of the points scales with credibility (credible interval range for reliable points in dark grey). The legend indicates the different orders and in parenthesis is the number of reaction norms per order.

The reaction norm shapes ranged from very steep to entirely flat (Fig. 4C), though steep reaction norms were more common than flat ones. The variance composition (variance among environments vs. variance among one’s offspring) correlated only very weakly with environmental predictability (Fig. 4C; 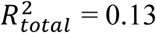, 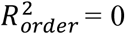, 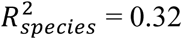, 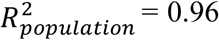, LRT ratio = 41.35, p < 0.0001), indicating little scope for the evolution of diversified bet-hedging. Similarly, we did not find any evidence for conservative bet-hedging, as the residuals in mean timing did not correlate with winter predictability (Fig. 4D; R^2^ = 0; LRT ratio = 0.45, p = 0.50).

The main results were robust to changes in the definition of winter onset, as a large range of temperature thresholds yielded similar R² values (Supplementary material S6). We also detected no temporal trend in reaction norm means or shapes with publication year (Supplementary Material S7).

## Discussion

### Mean timing

Insects and other invertebrates need to enter cold-resistant resting (diapause) stages before onset of winter. The timely induction of diapause is crucial for survival [36], so diapause reaction norms can be expected to be under intense selection pressure, and accordingly to be optimized to locally prevailing conditions. More than 50 years ago it has been reported that the critical day length, i.e. the inflection point of diapause reaction norms increases with latitude [34]. The latitudinal gradient was estimated as 60 – 90 minutes in day length change per 5°N, and this rule-of-thumb remains persistent in the literature [56–58]. Yet, even though the early empirical observations were based on few case studies with data of relatively low resolution, the findings have never been systematically validated. Our meta-analysis integrates data from 60 high-quality studies and applies robust statistical approaches. We did find a clear correlation of critical day length with latitude (Fig. 4A), but the slope was with 48.54 minutes per 5°N considerably lower than expected. This is in surprisingly close agreement with the observed critical day length gradient. Thus, we do not only provide strong empirical evidence for Danilevskii’s observation, but also provide a more reliable estimate and support it with climate data.

Importantly, the expected critical day length shift was not linear, but followed an exponential function (Fig. S4). With increasing latitude the onset of winter shifts linearly to earlier dates, but simultaneously autumn days are longer at high latitudes. To account for the complex relationship among day length, latitude and day of the year, we therefore converted the critical day lengths to ordinal days. The day of the year on which diapause would occur correlated reasonably well with the timing of mean winter onset at the lower latitudes (Fig. 4B), but less so for more northern populations. This is as expected from the observed linear critical day length change and the required exponential pattern. We need to emphasize, however, that the northern populations are putatively biased, as they are driven by data from a single genus (*Drosophila*) and region (Scandinavia, Fig. 3). More research is therefore needed to determine whether there is a real lack of diapause evolution in northern populations, but we demonstrate that shifts in critical day lengths are potentially not the main cues to which diapause is adaptively adjusted.

### Bet-hedging and plasticity

Demonstrating variance in a trait is not sufficient for diversified bet-hedging without showing that it increases geometric mean fitness in a population’s respective environment [59]. Correlating trait variance with environmental variance, indicates, however, whether bet-hedging is probable [59]. We here targeted reaction norms in four or more environments, thus allowing decomposing the variance in an among-environmental and a within-environmental component [25] that was subsequently correlated with among-year variability of winter onset. The lack of its strong correlation with variable climates did refute a strong signal for diversified bet-hedging (Fig. 4C). While we found that day length reaction norms are more variable than is commonly acknowledged, the correlation with environmental variability was low, which indicates that photoperiodic reaction norms are not generally used to hedge one’s evolutionary bets.

We can only speculate about the reasons for an apparent lack of bet-hedging. One reason may be the use of global threshold criteria that ignore interspecific variation in life history and cold tolerances (see limitations). Yet, mean diapause timing correlated well with mean winter onset, indicating that the criterion was a reasonable choice. We find it more likely that photoperiodic reaction norms have indeed not generally evolved as diversified bet-hedging strategies. Unpredictable conditions may, for instance, have instead selected for variance in life history strategies, such as variability in voltinism [60], in prolonged diapause over multiple years [61], or in mixed strategies of obligate and facultative diapause [54]. Alternatively, environmental conditions may be variable but predictable, e.g. when temperatures are temporally autocorrelated. Unfortunately, the thermal plasticity of reaction norms, voltinism and obligate diapause are rarely studied in conjunction with diapause reaction norms. We therefore call for further studies that shed light on the joint evolution of these strategies.

As alternative to diversified bet-hedging, unpredictable conditions may select for early diapause, so that the risk of fitness loss by early frost is mitigated at the cost of population growth [22,23] (conservative bet-hedging). We did, however, find neither a correlation between residual variation in mean phenology and environmental predictability, such that populations in highly unpredictable environments would diapause earlier than expected based on mean winter onset (Fig. 4D). Likely, an unknown species-specific time lag between diapause induction and winter onset severely limits our power to detect conservative bet-hedging.

### Evolutionary potential in a changing climate

Shifts in phenology play a key role in adapting to climate change [4,10,13], but there are concerns that constraints limit the evolutionary potential of phenology shifts. The current rate of change in climatic conditions is unprecedented [1], so predictions about future changes in evolutionary strategies are difficult. However, patterns of past adaptation (e.g. after range expansion) may provide information about evolutionary constraints, and thus show the upper limit of evolvable strategies. Using such a correlative approach, we have shown that the mean diapause timing generally evolved to match local conditions. However, our climate data showed that extreme shifts of the day length reaction norms would be required at high latitudes, indicating that further reaction norm evolution under climate change will be constrained at high latitudes. The lack of data in northern regions prevents a full evaluation, but it appears that diapause reaction norms of northern populations indeed did not match environmental conditions. We could further find no signs of ongoing adaptation to accelerating climate change, as diapause timing in the studies was independent of publication year. It would be premature to arrive at a clear conclusion and further research on these northern populations is clearly warranted, but this apparent lack of diapause evolution is worrying in a changing climate: the discrepancy between optimal and achieved diapause timing would continue to increase as species shift their range northwards, which may increase the extinction risk at the already vulnerable [62] northern edge of species distributions. Of course, northern populations may evolve alternative strategies (e.g. thermal plasticity, cold acclimation), but the evolutionary potential is also not known and warrants further exploration.

Genetic adaptation of the mean is not the only viable strategy in a changing climate. Projected increases in climate variability [21] can be expected to select for a shift from plasticity to bet-hedging strategies, but apart from a few case studies [63] it is not known whether bet-hedging can readily evolve. We have shown that plastic reactions by developmental switches are common, even when bet-hedging would be expected. In the majority of cases, the reaction norms were very steep and thus lead to rapid change of phenotypes within a short time window. Such steep developmental reaction norms might lead to an evolutionary trap, unless they are accompanied by plasticity to other cues [19] or generalized phenotypic responses such as adaptations to cope with stress [20]. Based on past patterns of adaptation, it appears that the evolution of flat (but not canalized) reaction norms is indeed constrained, leaving species vulnerable to changes in climate variability, unless bet-hedging can be achieved by other means.

### Data limitations and future research directions

A meta-analysis is naturally limited by the available data. We wish to summarize key areas in which our analysis remains speculative, and in which further data is crucial to arrive at a better understanding of reaction norm evolution.

First, we had to assume a common threshold of winter onset (fifth day of the year with temperatures below 10°C) across invertebrate orders. Future studies on the joint evolution of cold tolerance and critical day lengths will be invaluable to determine the adaptive evolution of reaction norm means in a changing climate. Secondly, the phylogenetic clustering made it impossible to account for differences in life history, which is strongly correlated with phylogeny. The life history affects the time between cue sensing and diapause induction, and thereby may interfere with a correct attribution of conservative bet-hedging. We hence call for more studies on currently understudied phyla. Third, we noted a lack of strongly canalized reaction norms. For butterflies it is well established that northern populations have frequently fewer generations per year, up to univoltine patterns with obligate diapause (‘sawtooth’ patterns [60]), but we found only few reaction norms that were flat at 0 or 100% diapause induction. We suspect that the lack of canalized reaction norms reflects study or reporting bias, because studies on diapause timing are unlikely to be conducted on obligately diapausing populations. The joint evolution of life cycles and critical day lengths (and bet-hedging therein) at high latitudes warrants further exploration. Lastly, we focused solely on day length reaction norms. Day length is widely acknowledged as the most important cue to time diapause [29,30,34], but warm temperature can in many organism delay the induction of diapause. It would hence be fruitful to study how day length reaction norms are modulated by temperature.

Taken together, we identified a crucial need for studies that study the joint effects of day length and temperature plasticity, integrate cold acclimation, and focus on currently understudied regions and species. Especially the populations at high latitudes warrant further study, because here the evolution of alternative strategies is most probable, while the data so far is also most scarce and phylogenetically and geographically clustered.

## Statement of authorship

JJ and DB designed the study; JJ performed the meta-analysis and wrote the initial draft of the manuscript; all authors were involved in interpretation of results and revisions of the manuscript.

## Data accessibility statement

Data and analysis scripts are available online: http://dx.doi.org/10.5061/dryad.9kd51c5d1

## Competing interest disclosure

The authors of this preprint declare that they have no financial conflict of interest with the content of this article.

An earlier version of this preprint (version 3) has been peer-reviewed and recommended by Peer Community In Ecology (https://doi.org/10.24072/pci.ecology.100040).

## Acknowledgements

This research has benefitted from a statistical consult with Ghent University FIRE (Fostering Innovative Research based on Evidence). We thank the Bradshaw & Holzapfel group (University of Oregon), Nobuaki Ichijo and Masahito T. Kimura (Hokkaido University Museum) for going to great lengths to share their data. In addition we would like to thank Jan Baert and Thomas Hovestadt for discussion of this manuscript. JJ was financially supported by a DFG research fellowship. DB is funded by FWO project G018017N.

An earlier version of this preprint has been peer-reviewed and recommended by Peer Community In Ecology (https://doi.org/10.24072/pci.ecology.100040). We are thankful for four additional anonymous reviews that led us to rethink our interpretation of the results.

## Supporting information

The following supplementary material is available for this article:

**Supplementary material S1**: Overview of studies from which reaction norms were extracted.

**Supplementary material S2**: Search terms for meta-analysis.

**Supplementary material S3**: Details on the MCMC approach.

**Supplementary material S4**: Correlation of day length at winter onset with latitude. Grey line: linear prediction between 21 and 69°N, grey points = points outside this prediction.

**Supplementary material S5. Correlation of mean diapause timing with latitude.** Each data point represents 1 reaction norm (425 in total), size of the points scales with credibility (credible interval range for reliable points in dark grey). The legend indicates the number of reaction norms per order.

**Supplementary material S6: Sensitivity of the meta-analysis to threshold choice**. The meta-analysis was repeated for parameter choices between 0 and 15. Panel A shows 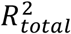 for the correlation of mean diapause timing with mean winter onset, panel B for the correlation of variance composition with day length predictability.

**Supplementary material S7**: Temporal trends in effect sizes across publication years. Black points indicate median of the MCMC estimate, grey lines show credible interval range.

## Notes

https://dx.doi.org/10.5061/dryad.9kd51c5d1

### Competing Interest Statement

The authors have declared no competing interest.

### Summary of Updates

After some constructive criticism of our interpretation we now discuss the limitations of our approach in more detail.

