## Supplementary Material for "Diapause is not selected as a bet-hedging strategy in insects: a meta-analysis of reaction norm shapes"

**Supplementary material S1**. Overview of studies from which reaction norms were extracted.

| Order | Genus | Species | Popula-tions | Reaction norms | Photo-periods | Region | Reference |
|---|---|---|---|---|---|---|---|
| **Coleoptera** | *Acanthoscelides* | *pallidipennis* | 3 | 3 | 5 | Japan | (Sadakiyo and Ishihara, 2011) |
| | *Bruchidius* | *dorsalis* | 3 | 3 | 5 | Japan | (Kurota and Shimada, 2003a) |
| | | *dorsalis* | 3 | 3 | 7 | Japan | (Kurota and Shimada, 2003b) |
| | *Harmonia* | *axyridis* | 4 | 4 | 5 | Europe, Asia | (Reznik et al., 2015) |
| | *Hippodamia* | *convergens* | 3 | 4 | 4 | US | (Obrycki et al., 2018) |
| | *Ips* | *typographus* | 4 | 4 | 5 | Europe | (Schroeder and Dalin, 2017) |
| | *Leptinotarsa* | *decemlineata* | 5 | 6 | 6 | Europe | (Lehmann et al., 2015) |
| | *Psacothea* | *hilaris* | 6 | 6 | 5 | Japan | (Shintani et al., 1996) |
| | | *hilaris* | 8 | 11 | 4 | Japan | (Shintani and Ishikawa, 1999) |
| **Diptera** | *Aedes* | *albopictus* | 21 | 21 | 12 | US, Japan | (Urbanski et al., 2012) |
| | | *atropalpus* | 3 | 3 | 5-7 | US | (Beach, 1978) |
| | | *sierrensis* | 5 | 5 | 4-7 | US | (Jordan and Bradshaw, 1978) |
| | | *triseriatus* | 8 | 8 | 10 | US | (Shroyer and Craig, 1983) |
| | *Boettcherisca* | *peregrina* | 6 | 6 | 8 | Japan | (Kurahashi and Ohtaki, 1989) |
| | *Chymomyza* | *costata* | 8 | 14 | 6-8 | Europe, Japan | (Riihimaa et al., 1996) |
| | *Drosophila* | *auraria* | 8 | 8 | 4-5 | Japan | (Kimura et al., 1993) |
| | | *auraria* | 7 | 7 | 4-7 | Japan | (Kimura, 1984) |
| | | *biauraria* | 11 | 11 | 4-5 | Japan | (Kimura et al., 1993) |
| | | *biauraria* | 4 | 4 | 5 | Japan | (Kimura, 1988) |
| | | *lacertosa* | 8 | 8 | 4-7 | Japan | (Ichijo, 1986) |
| | | *littoralis* | 8 | 8 | 7-11 | Europe | (Lumme and Oikarinen, 1977) |
| | | *littoralis* | 11 | 18 | 5-9 | Europe | (Lankinen, 1986) |
| | | *melanogaster* | 6 | 6 | 6 | Europe | (Pegoraro et al., 2017) |
| | | *montana* | 6 | 24 | 4-6 | Europe | (Tyukmaeva et al., 2011) |
| | | *subauraria* | 8 | 8 | 5-6 | Japan | (Kimura et al., 1993) |
| | | *subauraria* | 4 | 4 | 5-7 | Japan | (Kimura, 1984) |
| | | *takahashii* | 5 | 5 | 4 | Japan | (Kimura et al., 1994) |
| | | *triauraria* | 3 | 3 | 7-11 | Japan | (Yoshida and Kimura, 1994) |
| | | *triauraria* | 10 | 10 | 4-5 | Japan | (Kimura et al., 1993) |

| Order | Genus | Species | Popula-tions | Reaction norms | Photo-periods | Region | Reference |
|---|---|---|---|---|---|---|---|
| | | *triauraria* | 4 | 4 | 5-6 | Japan | (Kimura, 1984) |
| | *Sarcophaga* | *similis* | 4 | 4 | 7-8 | Japan | (Yamaguchi and Goto, 2019) |
| | *Wyeomyia* | *smithii* | 16 | 16 | 16-21 | US | (Bradshaw et al., 2003) |
| **Hemiptera** | *Laodelphax* | *striatellus* | 3 | 3 | 5-6 | Asia | (Hou et al., 2016) |
| | | *striatellus* | 8 | 8 | 6-8 | Japan | (Noda, 1992) |
| | *Nezara* | *viridula* | 5 | 5 | 5-8 | Japan | (Musolin et al., 2011) |
| | *Orius* | *sauteri* | 5 | 5 | 6-8 | Japan | (Shimizu and Kawasaki, 2001) |
| | | *sauteri* | 8 | 8 | 6-8 | Japan | (Ito and Nakata, 2000) |
| | *Rhopalosiphum* | *padi* | 3 | 3 | 11 | Europe | (Lushai and Harrington, 1996) |
| **Hymenoptera** | *Asobara* | *japonica* | 9 | 9 | 5 | Japan | (Murata et al., 2013) |
| | *Nasonia* | *vitripennis* | 7 | 7 | 8 | Europe | (Paolucci et al., 2013) |
| **Lepidoptera** | *Atrophaneura* | *alcinous* | 6 | 6 | 5 | Japan | (Kato, 2005) |
| | *Diatraea* | *grandiosella* | 3 | 3 | 6 | US | (Takeda and Chippendale, 1982) |
| | *Helicoverpa* | *armigera* | 5 | 5 | 6 | Asia | (Chen et al., 2013) |
| | | *armigera* | 3 | 3 | 6 | Japan | (Qureshi et al., 2000) |
| | | *armigera* | 3 | 3 | 4-5 | Japan | (Shimizu and Fujisaki, 2006) |
| | | *armigera* | 3 | 3 | 5 | Japan | (Shimizu and Fujisaki, 2002) |
| | *Hyphantria* | *cunea* | 4 | 4 | 4-5 | Japan | (Gomi and Takeda, 1996) |
| | | *cunea* | 3 | 3 | 4-6 | Japan | (Gomi et al., 2009) |
| | *Inachis* | *io* | 3 | 3 | 9 | Europe | (Pullin, 1986) |
| | *Leucoma* | *candida* | 5 | 5 | 4-5 | Japan | (Kuwana, 1986) |
| | *Papilio* | *glaucus* | 3 | 3 | 8-11 | US | (Ryan et al., 2018) |
| | | *memnon* | 4 | 4 | 8 | Japan | (Yoshio and Ishii, 1998) |
| | *Pararge* | *Aegeria* | 4 | 4 | 4 | Europe | (Lindestad et al., 2019) |
| | *Phyllonorycter* | *ringoniella* | 5 | 5 | 4 | Japan | (Ujiye, 1985) |
| | *Pieris* | *rapae* | 7 | 7 | 5-8 | Japan | (Hashimoto et al., 2008) |
| | *Sericinus* | *montelus* | 6 | 6 | 9 | Asia | (Wang et al., 2012) |
| | *Zygaena* | *trifolii* | 5 | 5 | 8-11 | Europe | (Wipking, 1988) |
| **Neuroptera** | *Chrysopa* | *oculata* | 9 | 9 | 4-6 | US | (Nechols et al., 1987) |

| Order | Genus | Species | Popula-tions | Reaction norms | Photo-periods | Region | Reference |
|---|---|---|---|---|---|---|---|
| **Thysanoptera** | *Haplothrips* | *brevitubus* | 3 | 13 | 6 | Japan | (Fujimoto et al., 2014) |
| | *Thrips* | *nigropilosus* | 6 | 6 | 4-6 | Japan | (Nakao, 2011) |
| **Trombidiformes** | *Tetranychus* | *pueraricola* | 33 | 33 | 5 | Japan | (Suwa and Gotoh, 2006) |
| | | *urticae* | 10 | 10 | 7-9 | Europe | (Vaz Nunes et al., 1990) |
| | | *urticae* | 5 | 5 | 6-8 | Japan | (So and Takafuji, 1992) |
| | | *urticae* | 8 | 8 | 7-12 | Europe | (Koveos et al., 1993) |
| | | *urticae* | 6 | 6 | 5-11 | Japan | (Gotoh and Shinkaji, 1981) |

**Supplementary material S2**: Search terms for meta-analysis.

**Overview**

This document describes the second search in Web of Science on 10.12.2018.

We searched for

TS = (("day length" OR photoperiod* OR diapaus* OR hibern* OR dorman*) AND (geogr* OR "range" OR latitud* OR longitud* OR cline$ OR clinal))

but excluded all articles that were found in the first search, as well as all review articles, retractions and corrections. We then filtered the 6575 results by research area and invertebrate-related terms. In particular, we:

1) Included all entomological articles (634)
2) Included all articles with invertebrate taxa (75 terms) named in title, keywords or abstract (901 articles)
3) All zoological articles that name no vertebrate (61 terms) in the title (192 articles)
4) All articles from ecology, evolutionary biology and genetics which name no vertebrate, plant or microbe (80 terms) in their title (572 articles).
5) All articles from relevant other topics (11 topics) that name no human psychological condition, vertebrate, plant or microbe (85 terms) in their title (288)

From these 2562 articles we excluded all references that name aquatic environments, unless they also named terrestrial environments. 2528 articles remained.

**Details**

Below are the exact search terms, with search ID and number of hits in red.

#1 TS = ((photoperiodic AND (geogr* OR range)) OR (photoperiod* AND latitud*) OR (photoperiod* AND longitud*)) 1820

#2 (TS = (("day length" OR photoperiod* OR diapaus* OR hibern* OR dorman* ) AND (geogr* OR "range" OR latitud* OR longitud* OR cline$ OR clinal)) not #1) *AND* **DOCUMENT TYPES:** (Article OR Abstract of Published Item OR Book OR Book Chapter OR Data Paper OR Database Review OR Discussion OR Editorial Material OR Excerpt OR Letter OR Note OR Proceedings Paper) 6575

#3 #2 and SU = "entomology" 634

#4 #2 not #3 AND TS =(invertebrat* OR worm* OR annelid* OR platyhelminth* OR nematod* OR mollusc* OR gastropod* OR slug* OR snail* OR arthropod* OR chelicer* OR arachnid* OR aranea* OR acari OR tetranych* OR ixod* OR opilion* OR spider* OR *scorpio* OR tick$ OR mite$ OR harvestmen OR crustace* OR malostraca* OR isopod* OR woodlice OR oniscid* OR armadillium OR myriapod* OR chilopod* OR diplopod* OR pauropod* OR symphyla OR millipede* OR centipede* OR hexapod* OR collembol* OR springtail* OR insect$ OR blattodea OR *ptera OR mantodea OR odonata OR phasmatodea OR psocodea OR thysanura OR zygentoma OR psyllid* OR stenorrhyn* OR cockroach* OR beetle$ OR earwig* OR *fly OR *flies OR droso* OR mosquit* OR *bug$ OR aphid* OR adelgid* OR phyllox* OR *wasp$ OR (*bee OR *bees) OR (ant OR ants) OR mantis

OR grasshopper* OR locust* OR cricket* OR louse OR lice OR flea$ OR moth$ OR thrip* OR silverfish ) NOT TI = (paleo* or $chiroptera*) 901

#5 #2 not #3 not #4 AND SU = "Zoology" NOT TI =( palaeo* OR $vertebra* OR *fish* OR $amphib* OR $salientia* OR $anura* OR $caudata OR $salamand* OR newt$ OR $gymnophion* OR frog$ OR tadpole$ OR toad$ OR $reptil* OR $crocodil* OR *sauria* OR $squamat* OR $lizard* OR $lacert* OR $gekko* OR $serpent* OR $snake* OR $testudin* OR $turtle* OR $tortois* OR $mammal* OR $rodent* OR $sciurid* OR $hamster* OR *mouse* OR *mice* OR $squirrel* OR $rabbit* OR $hare OR $hares OR $chiropt* OR $bat OR $bats OR $myotis OR $sorciomorpha OR $soricid* OR $talpid* OR $shrew* OR $marmot* OR $mole OR $moles OR $primat* OR $carnivora OR $ursid* OR $ursus OR $felid OR $felids OR "$sea lion" OR "$fur seal" OR "$elephant seal" OR $marsupi* OR $goat* OR $sheep* OR $deer OR $cattle OR estrus OR suprachiasm*) 192

#6 #2 not #3 not #4 AND SU = (ENVIRONMENTAL SCIENCES ECOLOGY OR EVOLUTIONARY BIOLOGY OR GENETICS HEREDITY OR BIODIVERSITY CONSERVATION OR SOIL SCIENCE NOT Zoology) NOT TI = ( palaeo* OR $vertebra* OR *fish* OR $amphib* OR $salientia* OR $anura* OR $caudata OR $salamand* OR newt$ OR $gymnophion* OR frog$ OR tadpole$ OR toad$ OR $reptil* OR $crocodil* OR *sauria* OR $squamat* OR $lizard* OR $lacert* OR $gekko* OR $serpent* OR $snake* OR $testudin* OR $turtle* OR $tortois* OR $mammal* OR $rodent* OR $sciurid* OR $hamster* OR *mouse* OR *mice* OR $squirrel* OR $rabbit* OR $hare OR $hares OR $chiropt* OR $bat OR $bats OR $myotis OR $sorciomorpha OR $soricid* OR $talpid* OR $shrew* OR $marmot* OR $mole OR $moles OR $primat* OR $carnivora OR $ursid* OR $ursus OR $felid OR $felids OR "$sea lion" OR "$fur seal" OR "$elephant seal" OR $marsupi* OR $goat* OR $sheep* OR $deer OR $cattle OR estrus OR suprachiasm*OR microb* OR bacteria* OR fung* OR *ceae OR bloom OR yield OR germination OR molecular OR simulation OR QTL OR spring OR cell* OR tiller OR cultivar* OR bud* OR chill* OR (tree NEAR phenology)) 572

#7 #2 not #3 not #4 not #5 not #6 NOT SU = (ENTOMOLOGY OR ZOOLOGY OR ENVIRONMENTAL SCIENCES ECOLOGY OR EVOLUTIONARY BIOLOGY OR GENETICS HEREDITY OR BIODIVERSITY CONSERVATION OR SOIL SCIENCE OR AGRICULTURE OR PLANT SCIENCES OR FORESTRY OR FOOD SCIENCE TECHNOLOGY) AND SU =(SCIENCE TECHNOLOGY OTHER TOPICS OR LIFE SCIENCES BIOMEDICINE OTHER TOPICS OR ENDOCRINOLOGY METABOLISM OR NEUROSCIENCES NEUROLOGY OR PHYSIOLOGY OR REPRODUCTIVE BIOLOGY OR INFECTIOUS DISEASES OR BEHAVIORAL SCIENCES OR ANATOMY MORPHOLOGY OR HEMATOLOGY OR HEALTH CARE SCIENCES SERVICES ) NOT TI = (human OR sleep* OR disorder OR depress* OR palaeo* OR $vertebra* OR *fish* OR $amphib* OR $salientia* OR $anura* OR $caudata OR $salamand* OR newt$ OR $gymnophion* OR frog$ OR tadpole$ OR toad$ OR $reptil* OR $crocodil* OR *sauria* OR $squamat* OR $lizard* OR $lacert* OR $gekko* OR $serpent* OR $snake* OR $testudin* OR $turtle* OR $tortois* OR $mammal* OR $rodent* OR $sciurid* OR $hamster* OR *mouse* OR *mice* OR $squirrel* OR $rabbit* OR $hare OR $hares OR $chiropt* OR $bat OR $bats OR $myotis OR $sorciomorpha OR $soricid* OR $talpid* OR $shrew* OR $marmot* OR $mole OR $moles OR $primat* OR

$carnivora OR $ursid* OR $ursus OR $felid OR $felids OR "$sea lion" OR "$fur seal" OR "$elephant seal" OR $marsupi* OR $goat* OR $sheep* OR $deer OR $cattle OR estrus OR suprachiasm*OR microb* OR bacteria* OR fung* OR *ceae OR bloom OR yield OR germination OR molecular OR simulation OR QTL* OR arabidopsis OR spring OR cell* OR tiller OR cultivar* OR bud* OR chill* OR (tree NEAR phenology)) 288

#8 (#3 or #4 or #5 or #6 or #7 AND TS = (terrest*) ) or (#3 or #4 or #5 or #6 or #7 not TS = (marine* OR aquat* OR limno* OR water )) 2528

We repeated the search in the Web of Science KCI (Korean Journal Database), but without article type restrictions, in the Russian Science Citation Index (excluding reviews), and in the SciELO Citation Index (excluding reviews):

| Search number | KCI | RCI | SciELO |
|---|---|---|---|
| 1 | 4 | 30 | 52 |
| 2 | 72 | 64 | 134 |
| 3 | 0 | 9 | 7 |
| 4 | 11 | 11 | 13 |
| 5 | 0 | 3 | 3 |
| 6 | 0 | 11 | 7 |
| 7 | 5 | 5 | 8 |
| **8** | **14** | **38** | **37** |

**Supplementary material S3**: Details on the MCMC approach.

We derived midpoints and variance composition from reaction norms, but the data was relatively scarce (on average seven data points per reaction norm). Hence standard non-linear regression techniques did not always yield reasonable estimates, for example the slope could not be estimated when there was only one data point present on the sloped part of the reaction norm. Nevertheless, the range of the possible parameter space can be estimated with Markov chain Monte Carlo methods. We thus estimated the 4-dimensional credible parameter space and calculated the variance components based on this parameter space.

*MCMC specifications*

We used rjags (Plummer 2018) to run Markov chain simulations on each of the 447 reaction norms. We ran 4 replicate chains with lengths of 11,000 iterations and discarded a burn-in of 1,000 iterations. We specified our model with (eq. 5), and consequently chose the binomial density function to estimate the likelihood. If specified in the primary study, we used the sample sizes of each day length treatment as number of trials, otherwise we used the global average of the study. For those studies that did not mention sample sizes, we used a global average of 100 trials for each of the data points. We implemented uninformative priors for all four parameters. These were:

$b$ ~ unif {-100, 100}
$c$ ~ unif {0, 1}
$d$ ~ unif {$c$, 1}
$e$ ~ unif {$D_{min}$, $D_{max}$}, with $D_{min}$ and $D_{max}$ being the range of applied day length treatments, converted in ordinal days.

The upper limit of the logit-function was constrained to be higher than the lower limit, because otherwise switching between the two equal solutions (positive slope, $d > c$ and negative slope, $c < d$) would render the chain unidentifiable. Despite the relative data scarcity, the four replicate Markov chains mixed well in nearly all cases, providing a well-defined frequency distribution. We repeated the analyses on the untransformed day length reaction norms to obtain a critical day length estimate that is comparable to those obtained in earlier studies.

The MCMC algorithms provided a 4-dimensional parameter space to define continuous reaction norms, and we calculated the variance components of those curves that fall within the credible intervals. To do so, we followed the trace of the MCMC algorithm. For each iteration step we sampled 1000 equally spaced day lengths around the proposed inflection point $e \pm 100$ days, and performed the variance calculations (eq. 1-5) on the proposed parameters $b$,$c$,$d$ and $e$. Following the logic of the MCMC algorithm, we reported the 0.025 and 0.975 quantiles of the resulting frequency distribution as credible intervals.

$$\sigma^2_{within} \stackrel{\text{def}}{=} \frac{\sum p_x * (1-p_x)}{n}$$ ;n = number of treatments (eq. 1)

$$\sigma^2_{among} \stackrel{\text{def}}{=} \frac{\sum (p_x - \overline{p_x})^2}{n-1}$$ (eq. 2)

$$r = \frac{\sigma^2_{among}}{\sigma^2_{among} + \sigma^2_{within}}$$ (eq. 3)

$$\sigma^2_P = \sigma^2_{among} + \sigma^2_{within}$$ (eq. 4)

$$p(x) = c + \frac{(d-c)}{1+\exp^{(b*(x-e))}}$$ (eq. 5)

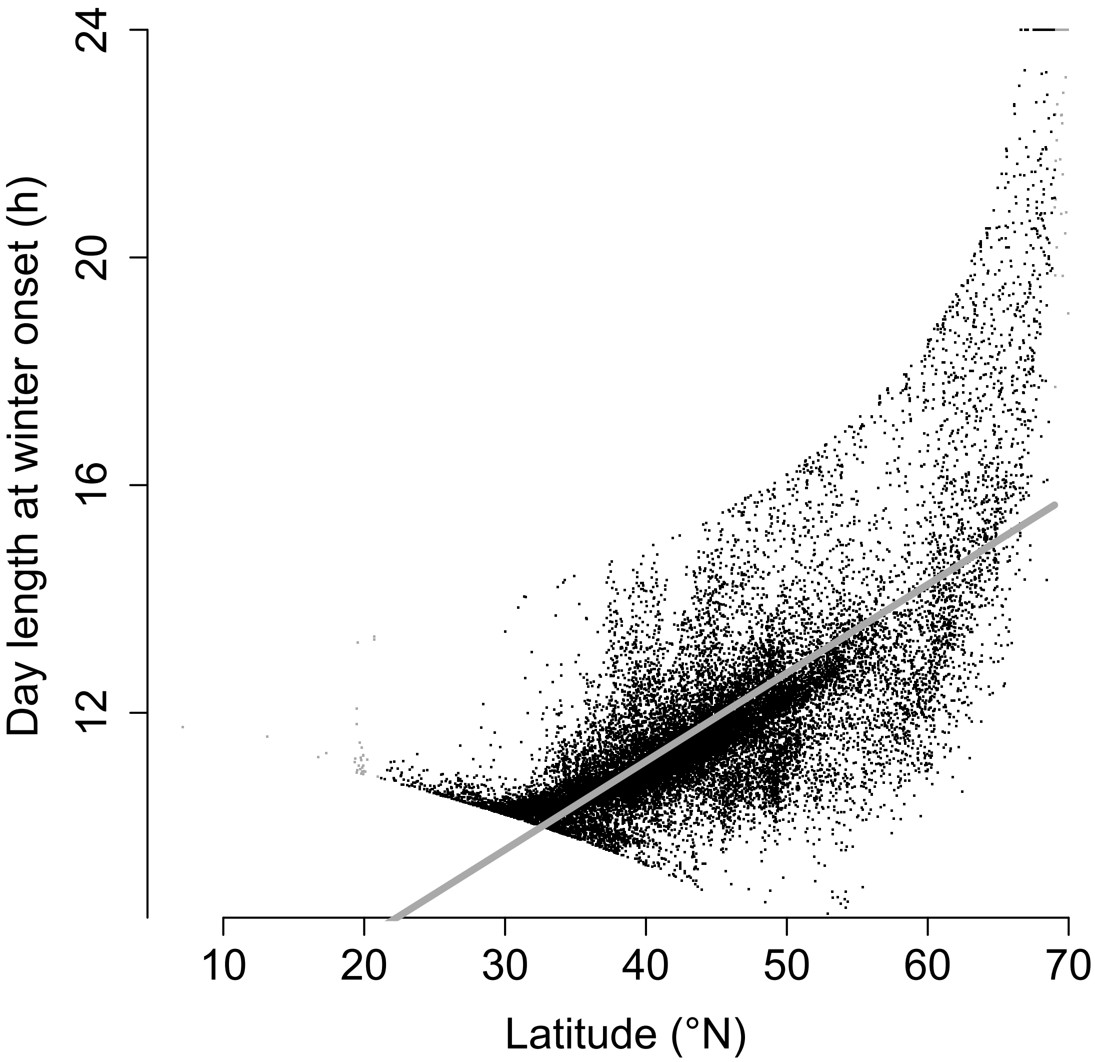


**Supplementary material S4**: Correlation of day length at winter onset with latitude. Grey line: linear prediction between 21 and 69°N, grey points = points outside this prediction.

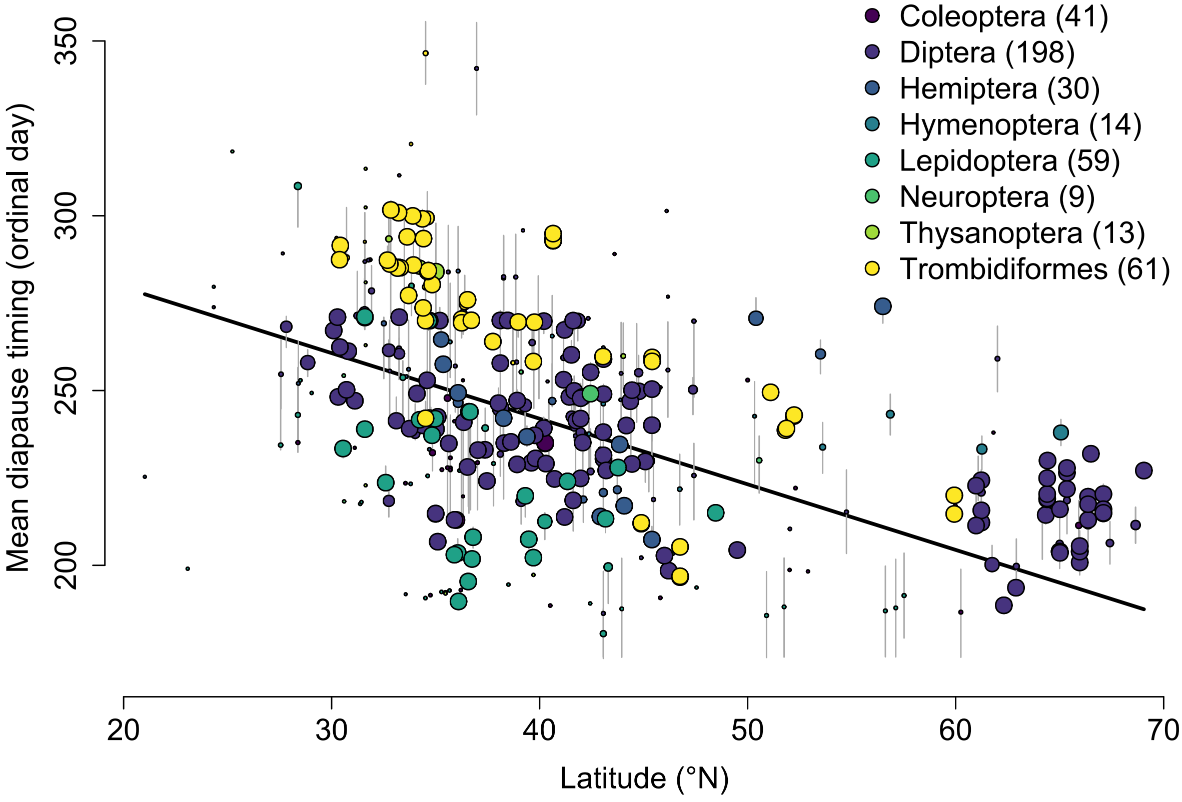


**Supplementary material S5. Correlation of mean diapause timing with latitude.** Each data point represents 1 reaction norm (425 in total), size of the points scales with credibility (credible interval range for reliable points in dark grey). The legend indicates the number of reaction norms per order.

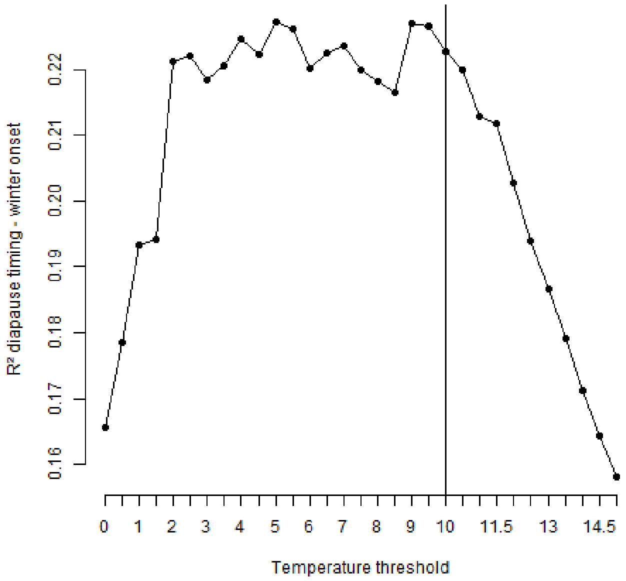


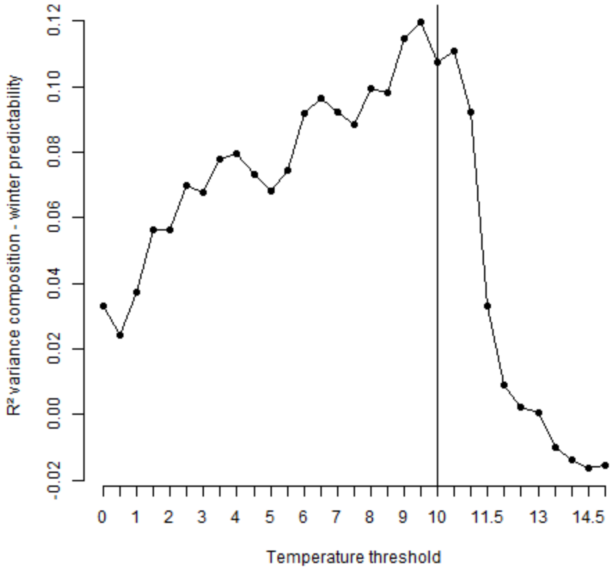


**Supplementary material S6: Sensitivity of the meta-analysis to threshold choice**. The meta-analysis was repeated for parameter choices between 0 and 15. Panel A shows $R^2_{total}$ for model 1 (Mean diapause timing vs. mean winter onset), panel B for model 2 (variance composition vs. day length predictability).

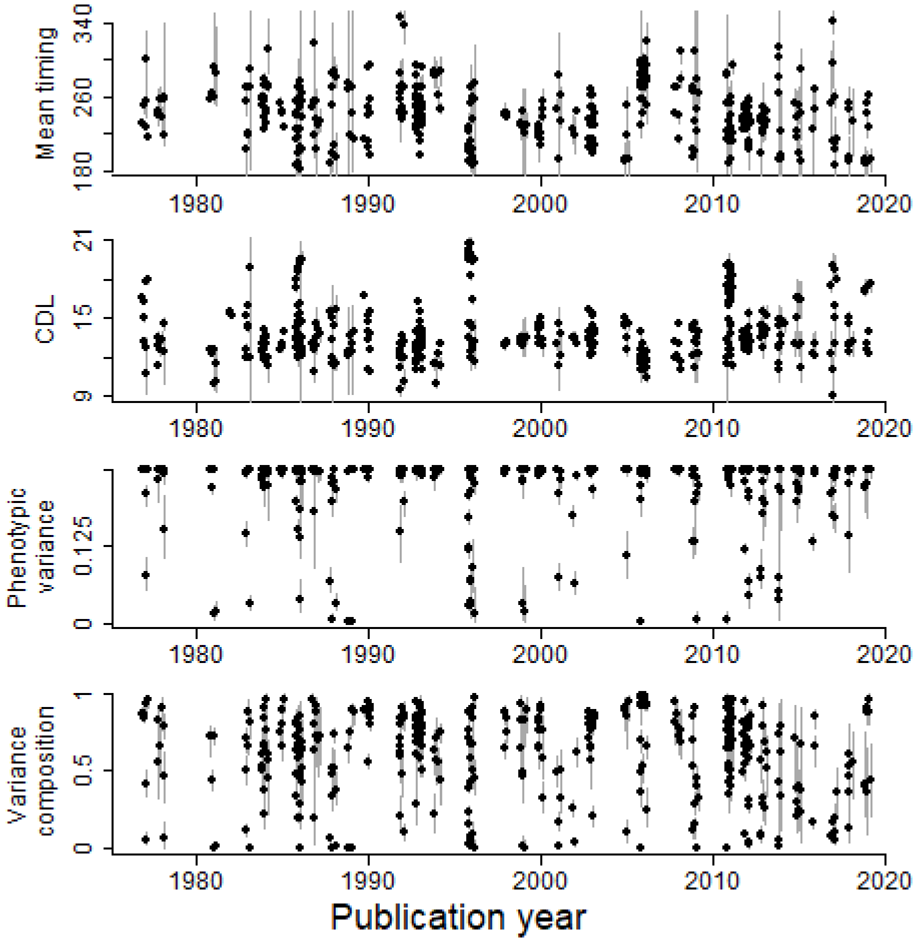


**Supplementary material S7:** Temporal trends in effect sizes across publication years. Black points indicate median of the MCMC estimate, grey lines show credible interval range.
